# Translational Regulation Promotes Oxidative Stress Resistance in the Human Fungal Pathogen *Cryptococcus neoformans*

**DOI:** 10.1101/735225

**Authors:** Jay Leipheimer, Amanda L. M. Bloom, Christopher S. Campomizzi, Yana Salei, John C. Panepinto

## Abstract

*Cryptococcus neoformans* is one of the few environmental fungi that can survive within a mammalian host and cause disease. Although many of the factors responsible for establishing virulence have been recognized, how they are expressed in response to certain host derived cellular stresses is rarely addressed. Here we characterize the temporal translational response of *C. neoformans* to oxidative stress. We find that translation is largely inhibited through the phosphorylation of the critical initiation factor eIF2α by a sole kinase. Preventing eIF2α mediated translational suppression resulted in growth sensitivity to hydrogen peroxide (H_2_O_2_). Our work suggests that translational repression in response to H_2_O_2_ partly facilitates oxidative stress adaptation by accelerating the decay of abundant non-stress related transcripts while facilitating the proper expression of critical oxidative stress response factors. Carbon starvation, which seems to induce translational suppression that is independent eIF2α, partly restored transcript decay and the expression of the critical oxidative stress response transcript Thioredoxin Reductase 1 (T*RR1*). Our results illustrate translational suppression as a key determinant of select mRNA decay, gene expression, and subsequent survival in response to oxidative stress.

**Importance:** Fungal survival in a mammalian host requires the coordinated expression and downregulation of a large cohort of genes in response to cellular stresses. Initial infection with *C. neoformans* occurs at the lungs, where it interacts with host macrophages. Surviving macrophage derived cellular stresses, such as the production of reactive oxygen and nitrogen species, is believed to promote dissemination into the central nervous system. Therefore, investigating how an oxidative stress resistant phenotype is brought about in *C. neoformans* furthers our understanding of not only fungal pathogenesis but also unveils mechanisms of stress induced gene reprogramming. We discovered that H_2_O_2_ derived oxidative stress resulted in severe translational suppression and that this suppression was necessary for the accelerated decay and expression of tested transcripts. Surprisingly, compounding oxidative stress with carbon starvation resulted in a decrease in peroxide mediated killing, revealing unexpected synergy between stress responses.

## Introduction

*C. neoformans*, an encapsulated fungus that causes meningitis and respiratory infection in both immunocompetent and immunocompromised individuals, is estimated to affect 220,000 people annually (1). In the context of a human host, M1 macrophage activation has been found to be essential for fungal killing, which is believed to be mediated through the production of reactive oxygen and nitrogen species (ROS and RNS respectfully)(2–4). High levels of ROS can cause major disruptions in cellular functions through oxidation of proteins, lipids, and nucleic acids (5). Therefore, oxidants must be contended with quickly and the damage caused by them repaired. Subjecting *C. neoformans* cultures to hydrogen peroxide (H_2_O_2_), which generates ROS, has been found to induce the simultaneous transcriptional expression of stress response factors coupled with the downregulation of homeostatic mRNAs (6). In *Saccharomyces cerevisiae*, H_2_O_2_ is met with strong translational inhibition (7). This inhibition was found to be achieved partly through the suppression of active ternary complex, which affects the rate of translation initiation (8). In the cytoplasm, the 5’ ends mRNA possess a methylated guanosine (cap) that protects it from 5’-3’ exonucleases while the 3’ end is protected from 3’-5’ exonuclease activity by the presence of long tracts of adenines (poly-A tail) bound by Poly-A Binding protein (Pab1p)(9–12). The described elements that protect the mRNA from decay also promote the translation of the transcript. Therefore, under our current understanding it seems that the mRNA decay and translational machinery are competing for the same elements found on a transcript. Indeed, an inverse correlation has been found between an mRNA’s translation initiation rate and its half-life, suggesting that one predominates over the other under certain conditions (13).

For the first time, in *C. neoformans*, we have characterized the translational response to oxidative stress. We find that translation is severely inhibited in response to H_2_O_2_ in an eIF2α-dependent manner. We used puromycin incorporation and polysome profiling to show that oxidative stress does not result in complete translational inhibition and that many oxidative stress response mRNAs are able to associate with ribosomes. Our work supports translational inhibition as the driving force of initial oxidative stress induced decay of transcripts abundant in unstressed conditions, with eIF2α phosphorylation triggering the decay. Importantly, translational repression is a requirement for oxidative stress resistance and can be conferred by carbon starvation in and eIF2a-independent manner. Altogether, this work characterizes the interplay between mRNA translation and decay as it pertains to oxidative stress resistance in a human fungal pathogen.

## Methods

### Strains and Media

The strain of *Cryptococcus neoformans* used in these studies is a derivative of H99O that retains full virulence and melanization. C. neoformans was cultivated on YPD (1% yeast extract, 2% peptone, 2% dextrose) agar unless otherwise indicated. Cultures were grown and seeded at 30° C as previously described (14).

The *gcn2*Δ mutant strain was constructed as described previously (15). List of primers and the corresponding restriction enzymes sites are listed in **Table 1**. Gene deletion and complementation was confirmed by PCR and northern blot analysis. GCN2 complementation was also confirmed by western blot analysis against the phosphorylated form of eIF2α. 5’ UTR reporter constructs were assembled using NEBuilder^®^ HiFi DNA Assembly Cloning Kit (#E5520S New England Biolabs Ipswich, MA). All amplifications were carried out following the manufacturer’s guidelines. pBluescript containing the G418 resistance cassette was used as a vector in the assembly. Full and in-line construct incorporation into the vector was confirmed by sequencing with primers reading into the desired adjacent amplified region. G418 resistant colonies were selected following biolistic transformation and galactose induction of the mCherry fused reporter was confirmed by northern blot analysis probing for the mCherry ORF.

### Isolation of Whole Cell lysate

Whole-cell lysate from yeast cultures was obtained by glass bead((#9831 RPI) mediated mechanical disruption using Bullet Blender Gold (Next Advance Model:BB2U-AU) set to power level 12 for 5 minutes. Lysate for the purpose of western blot analysis was suspended in buffer containing15 mM HEPES [pH 7.4], 10 mM KCl, 5 mM MgCl2, 10 μl/ml HALT protease inhibitor [Thermo Scientific]. Mt Prospect, IL). Crude lysate was then centrifuged at 22,000 rcf at 4°C for 10 min. The cleared lysate was aliquot from cellular debris into a new tube. Lysate for the purpose of northern blot analysis was obtained similarly, with the exception of the buffer used. RLT (#79216 Qiagen) was used to inhibit RNA decay during lysis. RNA was then isolated from cleared lysate using the manufacture’s protocol (#74106 RNeasy Mini Kit Qiagen).

### Western Blot Analysis

Western blots were performed using a total of 25ug of total protein derived from lysate and suspended in Laemmli Sample Buffer (#1610737 BIO RAD). Proteins were separated by gel electrophoresis using BIO RAD Mini-Protean TGX Stain-Free 4-15% Tris Glycine Gels (# 4568085 BIO RAD). These gels are embedded with a reagent that fluoresces when bound to a protein. Following gel separation, the fluorescence was analyzed and quantified using the BIO RAD Gel Doc XR+ imager default settings to verify the equal loading of protein across samples. Nitrocellulose transfer was performed using the BIO RAD Trans-Blot Turbo and corresponding transfer stacks at the instruments default TGX setting (#170-4270RTA Transfer Kit BIO RAD). Immunoblotting was performed as previously described (16). Primary antibodies anti-EIF2S1 (ab3215 Abcam), Anti-mCherry (RABBIT) (#600-401-P16), and anti-puromycin 12D10 (#MABE343 Millipore) were applied at 1:1000 for 12-18 hours at 4°C.

### Stability Assays and Northern Blot Analysis

Mid-log-phase cells grown at 30°C in YPD or minimal media (YNB) supplemented with 2% dextrose till exponential phase for all experimental conditions involving lysate derived experimental approaches. Yeast cultures were subjected then subjected to 1mM of H_2_O_2_. At the same time, transcriptional inhibition was achieved by the addition of 1,10-phenanthroline (#131377 ALDRICH) (250 μg/ml), and cultures were returned to the incubator at 30°C. Experiments accessing steady state levels of mRNA were performed similarly, except the transcriptional inhibitor was not included. Experiments involving strains expressing 5’ UTR reporters were performed by growing cultures to mid-log in YNB with 2% dextrose, followed by pelleting and resuspension in YNB with 2% galactose to induce reporter transcription. The reporter transcription was then inhibited by the removal of galactose through washing with and resuspension in YNB with 2% dextrose with and without H_2_O_2_. Northern blot analysis was performed and imaged using 5 μg of RNA as previously described (17). The hybridized transcript signal was normalized to rRNA gel bands. The half-life of *RPL2* was determined by nonlinear regression of normalized *RPL2* over time (Graphpad).

### Polysome Profiling

Yeast were grown in a 2-liter baffled flask in YPD with shaking at 250 rpm at 30°C for 5 to 6 h, reaching an OD_600_ of ∼0.55 to 0.65. Polysome profiles were obtained as described previously (14). Yeast cells were then harvested in the presence of 0.1 mg/ml cycloheximide (#66-81-9 Acros Organic) and pelleted immediately at 3,000 rcf for 2 min at 4°C. The yeast pellet was then flash frozen in liquid nitrogen, resuspended, and washed in polysome lysis buffer (20 mM Tris-HCl [pH 8], 2.5 mM MgCl, 200 mM KCl, 1 mg/ml heparin [#SRE0027-500KU], 1% Triton X-100, 0.1 mg/ml cycloheximide). Yeast cells were then lysed mechanically by glass bead disruption, resuspended in 500 μl of polysomal lysis buffer, and centrifuged for 10 min at 16,000 × g, 4°C to obtain the cytosolic portion of the lysate. Total RNA (250 μg) in a 250-μl total volume was layered on top of the polysome sucrose gradient (10% to 50% linear sucrose gradient, 20 mM Tris-HCl [pH 8], 2.5 mM MgCl, 200 mM KCl, 1 mg/ml heparin, 0.1 mg/ml cycloheximide). Gradients were subjected to ultracentrifugation at 39,000 rpm in an SW-41 rotor at 4°C for 2 h. Following centrifugation, sucrose gradients were pushed through a flow cell using a peristaltic pump, and RNA absorbance was recorded using Teledyne’s UA-6 UV-visible (UV-Vis) detector set at 254 nm. Absorption output was recorded using an external data acquisition device (DataQ). Fractions were then collected following absorption using a Teledyne retriever 500 set to collect 16-drop fractions.

To extract RNA, fractions were suspended in 3 volumes of 100% ethanol and incubated at −80°C for 12 to 16 h. The precipitate was collected via centrifugation at 16,000 × g at 4°C for 20 min and resuspended in 250 μl warm RNase-free water followed quickly with the addition of 750 μl Trizol LS (#10296010 Invitrogen). RNA was extracted per the manufacturer’s instruction. Purified RNA was resuspended in 30 μl RNase-free water. A third of this volume of each sample was used in subsequent Northern blot analyses.

### Puromycin Incorporation Assay

Yeast cultures were grown to mid-log for 5-6 hours in YNB supplemented with 2% dextrose. The main large culture of the experimental strains were then partitioned into separate containers and subjected to experimental conditions were indicated. 10 minutes before the indicated time point, 50ml volume of culture was taken then centrifuged and the resulting supernatant was removed from the yeast pellet. The pellet was then suspended in 5ml YNB 2% dextrose supplemented for 10 minutes with either 150 μg/ml puromycin (#P8833 Sigma), hydrogen peroxide, 100μg/ml cycloheximide, or a combination as indicated in the figure or figure legend. Short puromycin exposure time was used to limit the detrimental effects of aberrant protein build up that occurs due to the early termination of nascent polypeptide chains. After 10 minutes of puromycin incorporation lysate was acquired using the same method as described above.

### Flow Cytometry-Live/Dead Staining

Flow cytometry data was acquired using a BD LSRFortessa Cell Analyzer. Yeast were grown to exponential phase in minimal media supplemented with 2% dextrose. Cultures were then treated as outlined in figure 6D. After the two hour incubation step, cultures were washed with 1X PBS and suspended in 50μl 1X PBS. The Live-Dye Yeast Stain (#31062 Biotium Fremont, CA) protocol was performed as described by manufacturer, with the dead and thiazole orange stains incubated with cultures at room temperature for 30 minutes. Yeast were then fixed in a final concentration of 4% formaldehyde overnight at 4°. Samples were then diluted with a 3ml volume of 4% formaldehyde 1X PBS solution, which was used for sample input. FITC channel was used to detect thioazole orange, while the Texas red channel was used to detect dead yeast.

### Quantification and Statistical Analysis

Statistical analyses were performed using GraphPad Prism (version 6.05) software. Statistical analyses for stability data was obtained by determining the least squares fit of one-phase exponential decay nonlinear regression with Graphpad Prism software. Significance between curves was detected by sum-of-squares F test, with p < 0.05 determining that the data fall on separate regression lines, and therefore exhibit different rates of decay. Statistical analysis to compare mRNA abundance between Wild Type and the *gcn2*Δ mutant was performed using the Student’s t-test.

## Results

### Translation is temporally inhibited in response to oxidative stress in a dose dependent manner

In the ascomycete *S. cerevisiae*, translation is inhibited in response to the exogenous addition of H_2_O_2_ to the culture media and this inhibition is crucial for oxidative induced damage recovery (8). To determine the global translational state of the basidiomycete *C. neoformans* in response to oxidative damage, we chose to observe ribosome activity using two approaches. To compare the concentration of mRNA bound to free ribosomes, polysome profiles were derived from cultures grown to exponential phase at 30°C (unstressed) and after 30 minutes of exposure to 1mM H_2_O_2_. Polysome profiling, which examines the extent of ribosome association with mRNA in lysate by separating out large macromolecular complexes based on density in a sucrose gradient subjected to ultracentrifugation, suggests that the majority of the *C. neoformans* 40S and 60S subunits are engaged with mRNAs at exponential phase (Figure 1A). However, in response to H_2_O_2_ most of these ribosomes dissociate from mRNA and instead are found in the less dense portion of the gradient. A profile of this nature strongly suggests that translational output is low and that many of the free ribosome subunits are unable to associate with mRNA due to either a lack of active initiation factors or possibly a decrease in the total translatable mRNA.

**Figure 1.**
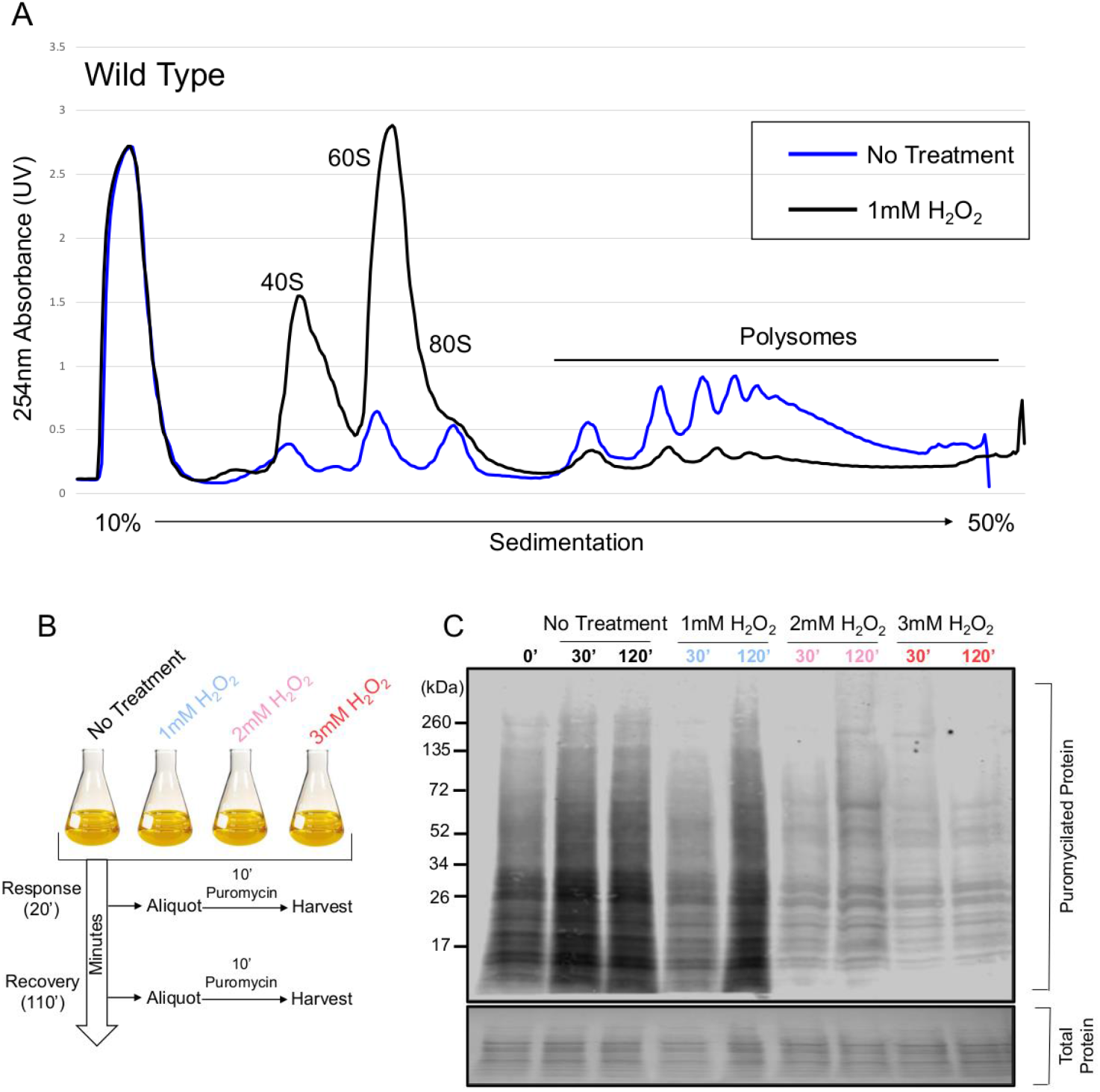
Translation is rapidly repressed in response to hydrogen peroxide induced oxidative stress. **(A)** Cultures were grown to exponential phase in YPD at 30°C before incubation in the presence or absence of 1mM H_2_O_2_ for 30 minutes. Absorption peaks derived from the lower portion of the gradient represent rRNA making up ribosomes bound to mRNA. Higher peaks in the heavy portion (polysomal) of the gradient compared to the lighter portion (Sub-polysomal) in the untreated condition indicate that most ribosomes at this point are engaged in translation. This pattern is reversed upon treatment with H_2_O_2_ with many of these ribosomes dissociating from mRNA and instead sediment with the sub-polysomal fraction. **(B)** Visual representation of experimental strategy for the assay shown in C. Cultures were subjected to varying concentrations of H_2_O_2_ with translational output assessed at two time points. **(C)** Visualization of puromycin incorporation in cultures exposed or not to varying concentrations of H_2_O_2_ at 30°C for either 30 or 120 minutes. Puromycin incorporation was detected by western blot with and anit-puromycin secondary antibody. Equivalent loading is indicated by the total protein stain.

Despite severe translational suppression in response to H_2_O_2_, a subset of mRNAs remain associated with ribosomes in the heavy polysome fraction. We, however, could not assume that these ribosomes are actively decoding the bound mRNA as prior studies report that translational elongation may be the target of inhibition in response to oxidative stress (18). Therefore, we probed the translational output of *C. neoformans* in response to varying concentrations of hydrogen peroxide, in vivo, using puromycin incorporation as a readout. Puromycin, which is an aminonucleoside antibiotic produced by the bacterium *Streptomyces albonigeris*, covalently binds to the growing nascent polypeptide chain during active translation (19). Higher overall rates of translational elongation and number of ribosomes engaged in elongation result in higher incorporation of puromycin, which can be detected in immunoblot assays using an antibody against the compound (**Supplementary Figure 1**). Because puromycin can be incorporated at any point along any transcript during elongation, the resulting molecular weights are dispersed throughout the western blot lane. Puromycin was added to the culture media 10 minutes prior to the indicated harvest time to limit any detrimental effects puromycin may have on growth (**Figure 1B**). Increasing concentrations of hydrogen peroxide resulted in like decreases in translational output at 30 minutes post H_2_O_2_ exposure (Figure 1C). Likewise, increasing concentrations of H_2_O_2_ extended the length of time cultures spent in a translationally repressed state. Therefore, these results indicate that *C. neoformans* is able to regulates the extent of translational inhibition in response to the severity of the oxidative stress. However, translation is not completely inhibited even in the presence of higher levels of oxidative stress suggesting that a subset of mRNAs and ribosomes are resistant to translational repression induced by H_2_O_2_.

### Oxidative stress induced translational repression is driven by the phosphorylation of elF2α and is required for oxidative stress resistance

We next aimed to discover the underlying mechanism, in *C. neoformans*, that facilitates such rapid and severe translational suppression in response to oxidative stress. Translation is partly regulated at the level of initiation, which is considered the rate limiting step of protein synthesis. A major molecular strategy used by eukaryotes in response to a variety of stresses is to limit the availability of functional initiator tRNA ternary complex, which is required for subunit joining at the canonical start codon (20). Phosphorylation of the α subunit within the elF2 complex at a conserved serine position prevents the recycling of new initiator tRNA following prior delivery to the start codon, thereby, preventing translation initiation through canonical means (21). To assess the status of eIF2α in response to oxidative stress in *C. neoformans*, we performed immunoblots using an antibody that recognizes the phosphorylated form of eIF2α. Increasing concentrations of hydrogen peroxide resulted in increased levels of overall phosphorylation of elF2α following a 30-minute exposure (**Figure 2A**). Whereas levels of elF2α phosphorylation return to basal levels in cultures exposed to lower concentration of H_2_O_2_ by two hours post-exposure, subjecting cultures with two and three millimolar concentrations resulted in sustained levels of eIF2α phosphorylation. The extent of eIF2α phosphorylation correlated well with the degree of translational repression seen in the above puromycin incorporation assays (**Figure 1C**).

**Figure 2.**
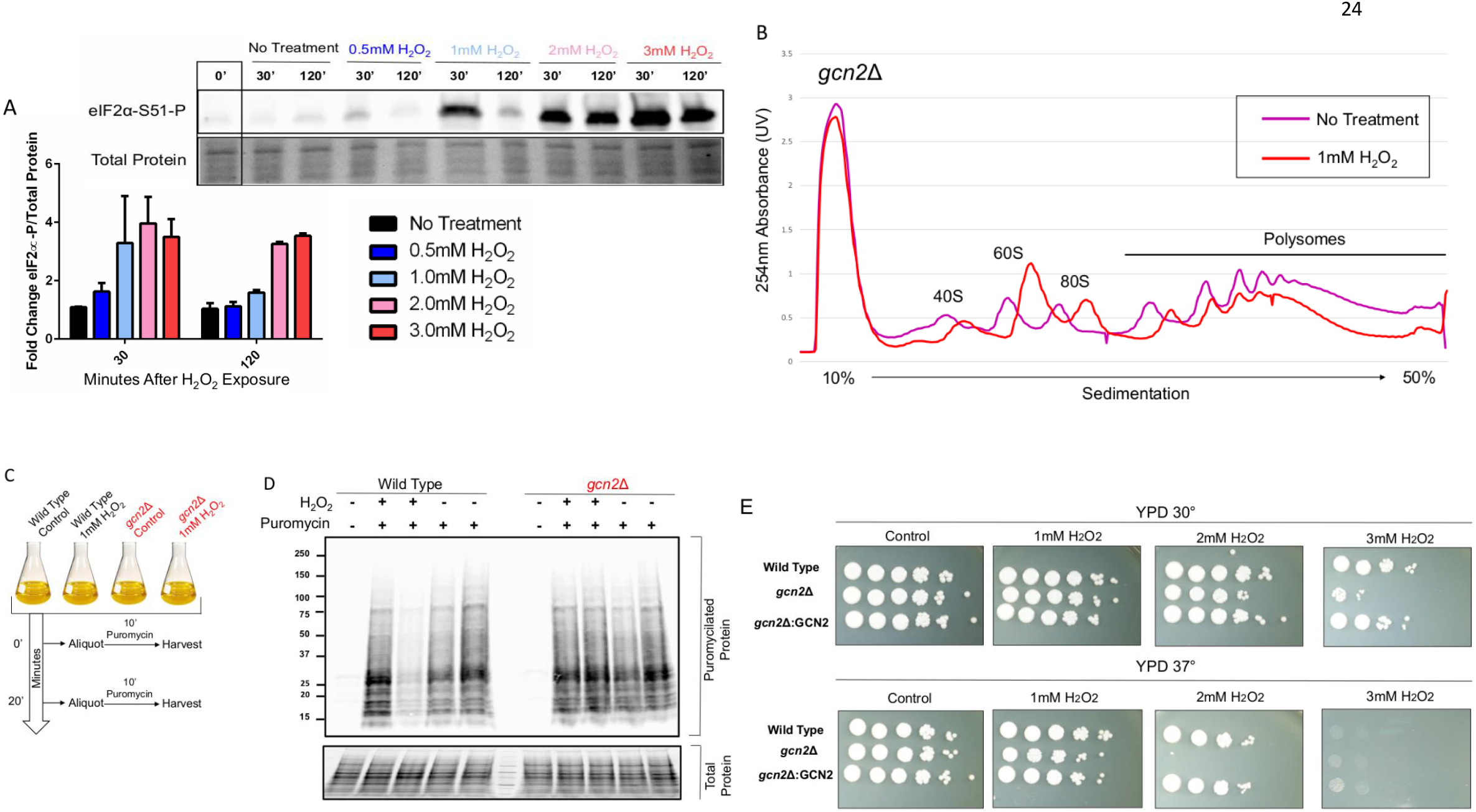
eIF2α is phosphorylated Gcn2 upon exposure to hydrogen peroxide in a dose dependent manner. **(A)** Phosphorylation status of eIF2α was assessed by western blot analysis using an antibody recognizing the phosphorylated form of the epitope. Cultures were grown to midlog in minimal media supplemented with 2% dextrose and were then treated with the indicated concentrations of H_2_O_2_. Cultures were harvested 30 and 120 minutes after treatment. Time point 0 represents cultures prior to exposure and was used to determine the fold change in phosphorylation. Equivalent loading is indicated by the total protein stain. n=3 **(B)** Cultures were grown to exponential phase in YPD at 30°C before incubation in the presence or absence of 1mM H_2_O_2_ for 30 minutes as done in figure 1A. n=4 **(C)** Schematic highlighting experimental treatment prior to lysis. Cultures were grown to midlog in minimal media supplemented with 2% dextrose before treatment with or without 1mM H_2_O_2_. Puromycin was added 10 minutes prior to harvest at indicated times **(D)** Puromycilated proteins are visible by western blot analysis using an antibody against the covalently bound antibiotic. No signal was detected in cultures that were not subjected to puromycin. Equivalent loading is indicated by the total protein stain **(E)** Serial dilution assays were performed by suspending cultures grown for 16 hours at 30° C to an O.D.600 of 1.0 and serially diluting them 10 fold in a 96 well plate. 5ul of the diluted yeast suspension was plan placed on YPD imbued with varying concentrations of H_2_O_2_. Plates were then incubated at either 30° C or 37 ° for 2-3 days.

To assess the importance of eIF2α phosphorylation in promoting translational repression in response to oxidative stress, we sought to eliminate eIF2α kinase activity in *C. neoformans*. Sequence comparisons of known eIF2α kinases suggested that *C. neoformans* may only possess one candidate eIF2α kinase with homology to Gcn2 which, of the eIF2α kinase family members, has the widest distribution among eukaryotes (22). Ablating the gene encoding for Gcn2 in *C. neoformans* (CNAG_06174) resulted in the absence of any observable elF2α phosphorylation signal after exposure to H_2_O_2_ (**Supplemental Figure 2**). Having observed that levels of elF2α phosphorylation tightly correlate with levels of translational repression and that Gcn2 is required for this phosphorylation, we next performed polysome profiling in the *gcn2*Δ strain (**Figure 2B**). Compared to profiles derived from Wild Type lysate, where the higher peaks corresponding to the polysome fraction (mRNAs bound to two or more ribosomes) were reduced in response to H_2_O_2_, there was little ribosome dissociation in the absence of Gcn2. These results strongly suggest that ROS induced translational inhibition in *C. neoformans* is largely triggered by eIF2α phosphorylation (**Supplementary Figure 3**). It should be noted, however, that a minor decrease in the polysome peaks suggest that eIF2α independent means of translational suppression are still active. Puromycin incorporation assays performed in the *gcn2*Δ strain suggest that the observed polysome peaks are active ribosomes that possess the ability to incorporate new tRNAs (**Figure 2C-D**).

To address the phenotypic consequence of preventing eIF2α phosphorylation in response to oxidative stress, we performed serial dilution assays using the Wild Type strain (H99), the *gcn2*Δ strain, and the complemented strain (**Figure 2E**). There were no observable growth sensitives in the absence of Gcn2 when grown on culture media alone. However, the presence of the oxidative stressors H_2_O_2_ (**Figure 2E**), T-BOOH, and Nitric Oxide (NO^−^) (**Supplementary figure 4**) resulted in a severe growth sensitivity in the *gcn2*Δ strain that was exacerbated by increased concentrations and incubation temperature. Together, these results indicate that Gcn2 is required for the phosphorylation of eIF2α and that the resulting translational repression in response to H_2_O_2_ promotes oxidative stress adaptation.

### Preventing elF2α phosphorylation following oxidative stress results in dysregulation of oxidative stress response transcript levels

Experiments performed in model yeast and other eukaryotes suggest that limiting ternary complexes through eIF2α phosphorylation can favor non-canonical translation of certain transcripts in response to stress, such as those possessing upstream Open Reading Frames (uORF) (21, 23–25). In *C. neoformans*, many of the ROS response transcripts are predicted to possess extensively structured 5’ UTR with uORF, such as the oxidative stress response transcript *ERG110*, that would prevent the recognition of the annotated ORF. We hypothesized that preventing eIF2α from being phosphorylation in response to oxidative stress would translationally disfavor *ERG110*. While testing this hypothesis we were surprised to find that the radioactive signal from the northern blots probed for the *ERG110* transcript was much higher in the *gcn2*Δ strain compared to that of Wild Type (**Figure 3A**). It was immediately evident that, in the absence of Gcn2, there is an overabundance of *ERG110* mRNA present compared to the Wild Type strain. Furthermore, the transcript remains detectable throughout the experimental time points in the *gcn2*Δ strain, whereas levels return to basal in Wild Type by the end of the time course. The presence of such a dramatic transcriptional dysregulation led us to probe for another oxidative stress response transcript, yet one that displayed sustained levels throughout the experimental time course. Thioredoxin Reductase (*TRR1*) is an essential gene in *C. neoformans* responsible for reducing thioredoxin peroxidase (*TSA1*) as well as reducing enzymes responsible for synthesizing basic cellular components required for DNA damage repair, such as ribonucleotide reductase (26, 27). *TRR1* is induced in response to H_2_O_2_ and levels remain above basal for two hours following stress exposure (**Figure 3B**). However, in the absence of Gcn2, levels of *TRR1* are well below what is observed in Wild Type and do not display signs of returning to basal 2 hours post-exposure. The defect in *TRR1* levels does not seem to be due to a defect in the expression of the transcript’s respective transcription factor, as *ATF1* levels are higher than expected in the *gcn2*Δ strain compared to Wild Type (28)(**Supplementary Figure 5A**). It is important to note that not all transcripts are dysregulated in the absence of Gcn2 as levels of *TSA1*, which acts to reduce H_2_O_2_ and indirectly promote the expression of *TRR1*, were found to be equivalent under time points tested (29, 30)(**Supplementary Figure 5B**).

**Figure 3.**
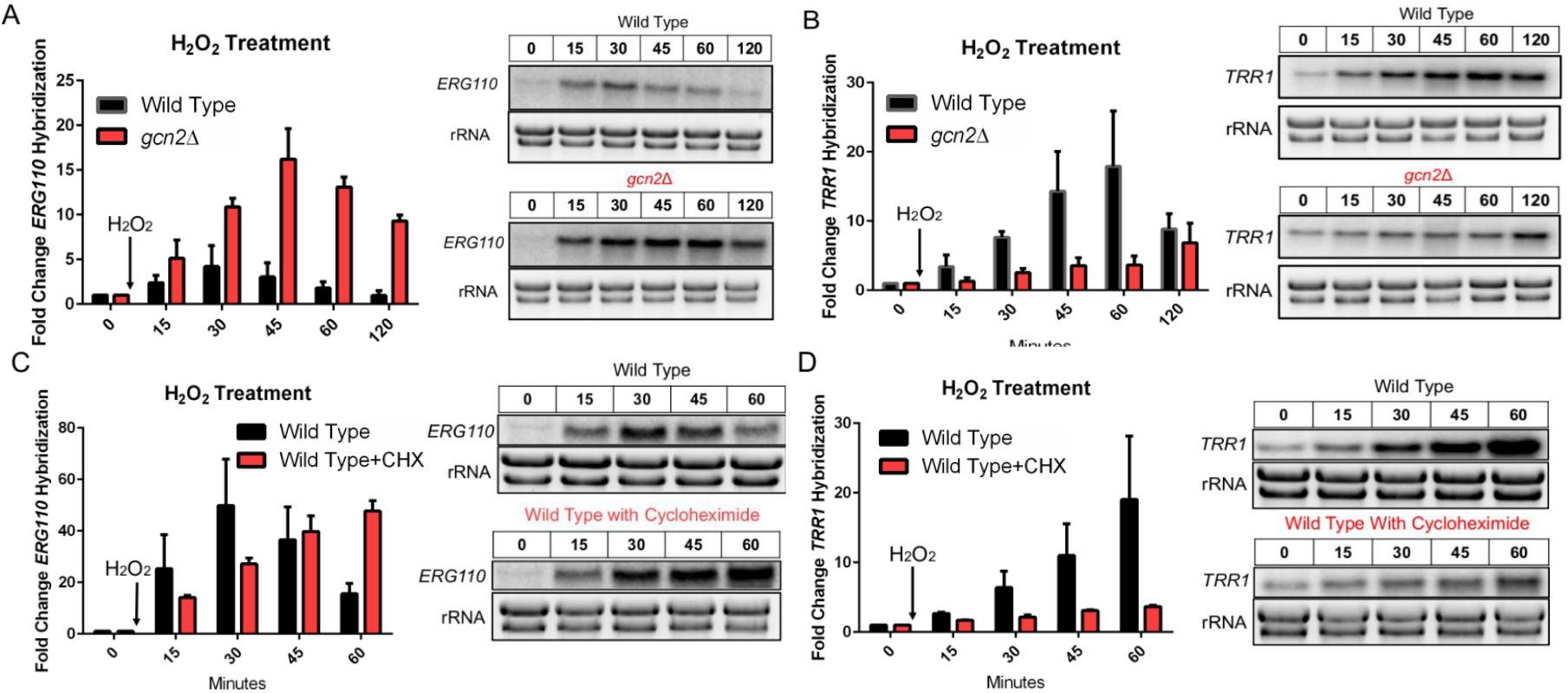
Gcn2 mediated translational inhibition facilitates the proper expression of oxidative stress response transcripts TRR1 and ERG110. Cultures were grown to exponential phase in YPD and where treated with 1mM H_2_O_2_. Aliquots were harvested at indicated time points and whole RNA was extracted. Northern blot analysis was performed probing for either **(A,C)** *ERG110* or **(B,D)** *TRR1*. The rRNA bands were imaged using SYBR Safe stain prior to membrane transfer and was used as both a loading control and to assess RNA sample integrity. (A-B) Steady state levels of *ERG110* (p=0.0185) and *TRR1* (p=0.0278) were assessed over a 2 hour time point following exposure to H_2_O_2_. Representative northern blot image is shown right of the corresponding bar graph. n=3 **(C-D)** Steady state levels of *ERG110* (p=0.0363) and *TRR1* (p=0.0199) were assessed over an 1 hour time point following exposure to H_2_O_2_ alone or with the addition of translation elongation inhibitor cycloheximide (CHX). n=2 A unpaired t test was used to determine if the mean differences between transcript levels of Wild Type and *gcn2*Δ were statistically significant. Error bars indicate SD between replicates.

Together, these results suggest that the absence of Gcn2 results in the dysregulation of certain stress response genes. To see if this dysregulation is related to the *gcn2*Δ strain’s inability to effectively clear mRNAs of ribosomes immediately following ROS derived stress, we repeated the prior experimental procedure only with the addition of the translation elongation inhibitor cycloheximide (**Figure 3C-D**). Preventing ribosome transcript run-off in response to hydrogen peroxide using cycloheximide in the Wild Type strain recapitulated the observed dysfunctional transcript levels observed in the *gcn2*Δ strain for both *ERG110* (**Figure 3C**) and *TRR1* (**Figure 3D**). These results suggest that polysomal collapse in response to oxidative stress seems to have an effect on the expression of oxidative stress response transcripts, which may stem from the availability of free ribosome subunits.

### Gcn2 is required for the accelerated decay of the “growth-related” transcript RPL2

Previous results in our lab have shown that many ribosomal protein (RP) transcripts undergo rapid decay in response to a variety of stresses (14, 31, 32). Having observed a defect in the expression of certain stress response transcripts, we asked if the accelerated decay of factors related to ribosome biogenesis in response to stress was also disrupted in the *gcn2*Δ strain. Northern blot analysis was performed probing for the Large Protein Ribosome Subunit 2 (*RPL2*) transcript following transcriptional shut-off and H_2_O_2_ exposure (**Figure 4A**). Where the half-life of *RPL2* is found to be around 40 minutes in Wild Type strain, the absence of Gcn2 resulted in a dramatic increase in stability with an unobtainable half-life under the observed time points. To see if the observed defect in *RPL2* transcript reduction was due to ribosomes remaining associated with mRNA in the *gcn2*Δ strain, we again isolated total RNA from the Wild Type strain following exposure to H_2_O_2_ and cycloheximide (**Figure 4B**). The drastic reduction in the levels of *RPL2* in response to H_2_O_2_ is completely mitigated by the simultaneous addition of the translation elongation inhibitor. These results suggest that, at least for our represented endogenous RP transcript, ribosome dissociation in response to peroxide stress may trigger the rapid decay of certain transcripts, and that peroxide stress leads to clearance of these mRNAs from the translational machinery.

**Figure 4.**
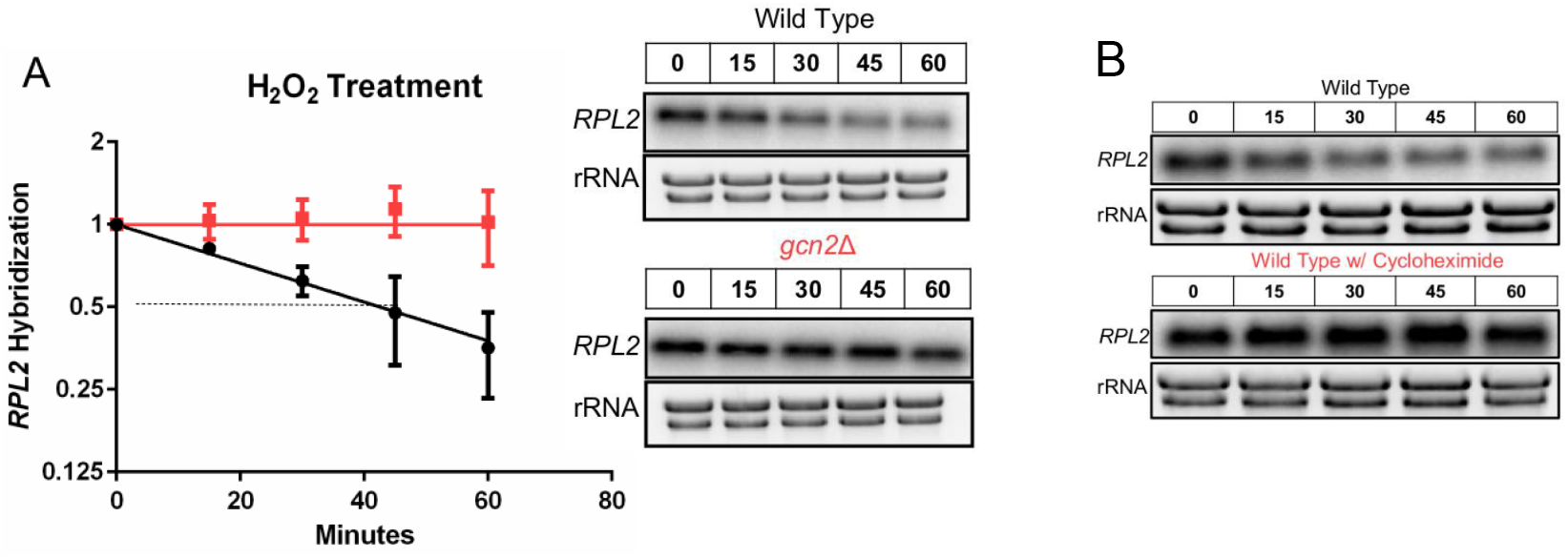
eIF2α phosphorylation is required for the accelerated decay of RPL2 in response to H_2_O_2_. Cultures were grown to exponential phase in YPD and were subjected to 1mM H_2_O_2_ in all experimental conditions. **(A)** 250ug/ml of 1-10 Phenanthroline was used at the start of the time course to inhibit transcription, allowing for the independent assessment of stability. Whole RNA was extracted and northern blot analysis was performed probing for *RPL2*. Representative northern blot image is shown right of the corresponding decay curve. n=3 **(B)** Representative northern blot image steady state levels of *RPL2* (p<0.0001) following exposure to 1mM H_2_O_2_ with or without the addition of the elongation inhibitor cycloheximide. n=3 Statistical analyses for stability data was obtained by determining the least squares fit of one-phase exponential decay nonlinear regression.

To determine if the increase in *RPL2* stability in the absence of Gcn2 was directly related to the translational state of that transcript, RNA was isolated from sucrose gradients following ultracentrifugation (**Figure 5A**). A transcript is considered highly translated if it is found more so in the high-density portion of the gradient, in contrast, a transcript is considered poorly translated if found distributed in the lower density portion. In unstressed conditions, the distribution of *RPL2* in both the Wild Type and the *gcn2*Δ strains is found in the higher density portion of the gradient (**Figure 5A Top Panel**). Where exposure to H_2_O_2_ in the Wild type strain resulted in the translational suppression of *RPL2* as observed by a shift in the distribution of the transcript to the lighter density portion of the gradient, the translational state of *RPL2* remained unchanged in the absence of Gcn2 (**Figure 5A Middle Panel-5B**). This further supports translational suppression as a method for accelerated decay in response to oxidative stress in *C. neoformans*.

**Figure 5.**
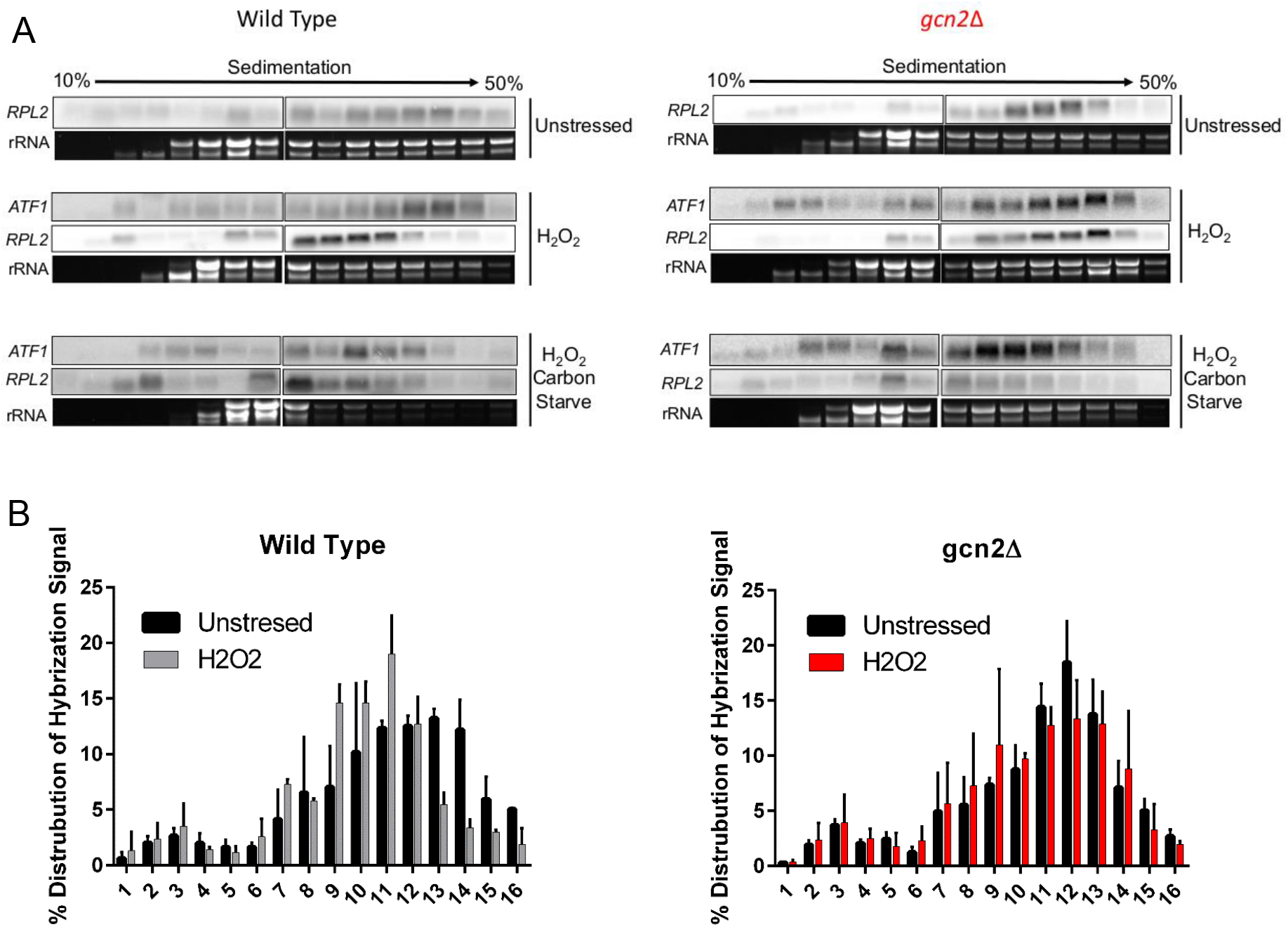
Gcn2 is required for the translational suppression of RPL2 in response to hydrogen peroxide but not carbon starvation. (A) RNA was isolated and precipitated from fractions acquired following polysome profiling as described in supplemental figure 3. All fractions were dissolved in the same volume of water and a third of that volume was used for northern blot analysis probing for either *RPL2* or *ATF1*. Prior to membrane transfer, rRNA was visualized and imaged to assess RNA stability and overall distribution though the differential sucrose gradient. Images represent results from three biological replicates (B) The hybridized signal intensity of *RPL2* in the Wild Type and gcn2Δ strain in unstressed and stressed conditions was determined for each fraction and summed. The intensity of each respective fraction compared to the total was use to quantify overall distribution of the signal throughout the gradient. Three biological replicates were performed. Error bars indicate SD.

**Figure 6.**
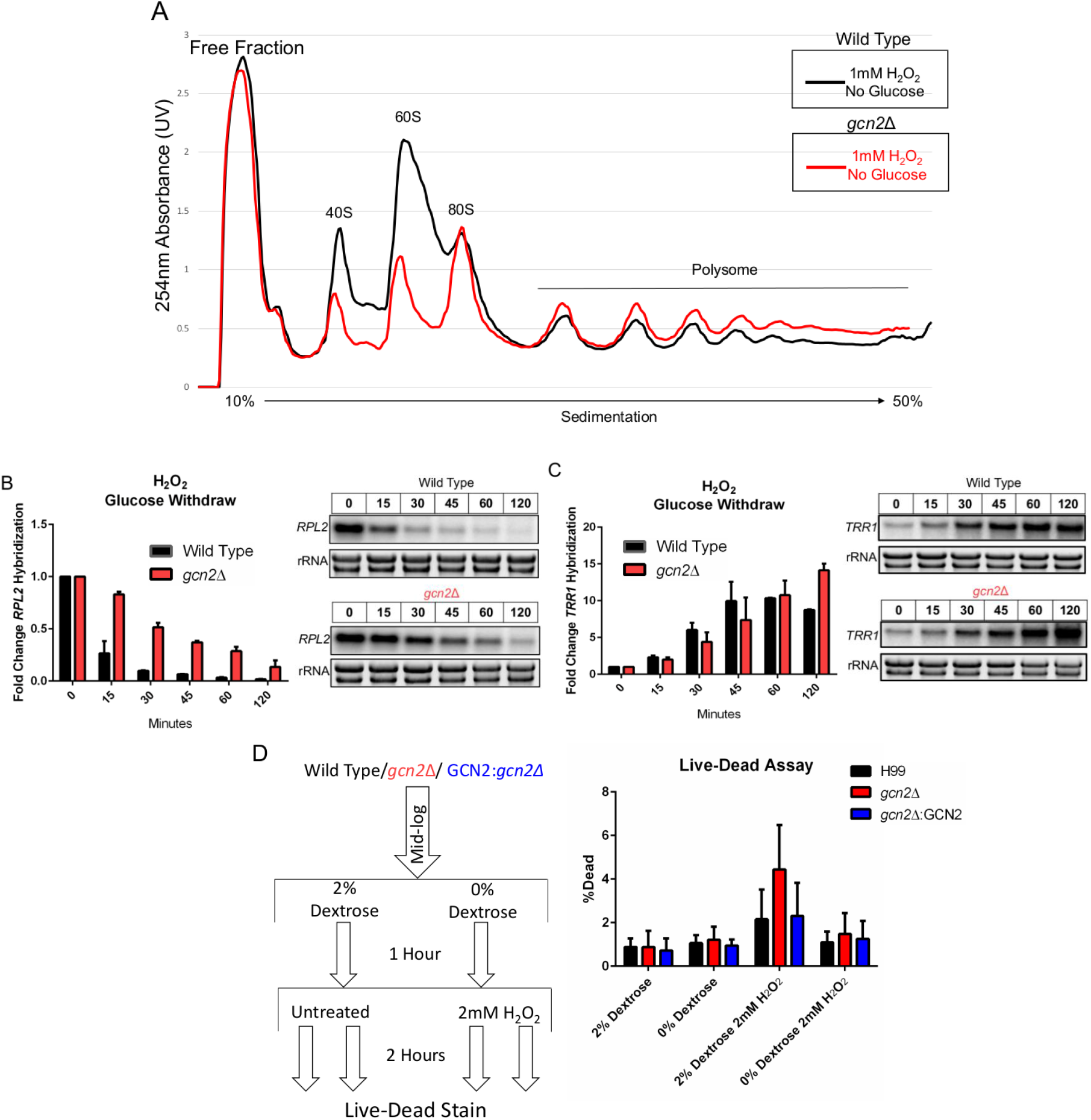
Glucose mediated translational repression is able to restore oxidative stress resistance in the absence of eIF2α phosphorylation. Cultures were grown to exponential phase in minimal media supplemented with 2% dextrose, at which point they were pelleted, washed with water, and suspended in minimal media alone supplemented with 1mM H_2_O_2_. **(A)** Polysome profiles were performed using yeast that were harvested and lysed 30 minutes after resuspension in carbonless minimal media lacking dextrose with 1mM H_2_O_2_. **(B-C)** At indicated time points following resuspension in carbonless minimal media containing 1mM H_2_O_2_, whole RNA was extracted and northern blot analysis probing for steady state levels of *RPL2* (p=0.0209) or *TRR1* (p=0.8497) was performed. Representative northern blot image is shown right of the corresponding bar graph (n=3). A paired t test was used to determine if the mean differences between transcript levels of Wild Type and gcn2Δ were statistically significant. Error bars indicate SD between replicates. **(D)** Yeast were grown to mid-log in minimal media supplemented with 2% dextrose. Cultures were then pelleted, washed in water, and re-suspended in minimal media with or without dextrose. After 1 hour, cultures were either subjected to 2mM H_2_O_2_ or left untreated for two hours prior to live-dead staining. The percentage of dead yeast were calculated from the total population, which was determined through the use of thiazole orange staining. n=3

### Glucose withdraw results in eIF2α-independent translational suppression and rescues ROS sensitivity in gcn2Δ strain

Having witnessed the consequences of continued ribosome association with mRNAs in response to H_2_O_2_, we wondered if we could rescue transcript expression and decay in the gcn2Δ strain by inducing eIF2α independent means of translational inhibition. Polysome profiles obtained in the *gcn2*Δ strain subjected to glucose withdraw in addition to 1mM H_2_O_2_ exposure, suggest that inhibition of translational in response to carbon starvation is still intact in the absence of elF2α phosphorylation (**Figure 6A-Supplemental Figure 3**). To further test our earlier hypothesis, that preventing ribosome association with certain mRNAs in response ROS is needed for both the proper removal and expression of transcripts, we performed a time course analysis of total RNA following the simultaneous removal of glucose and addition of H_2_O_2_. Carbon starvation partially restored the accelerated decline of *RPL2* in the *gcn2*Δ strain (**Figure 6B**). Carbon starvation also restored translational suppression of *RPL2* as observed in the disruption of the transcripts to the lower density portion of the gradient (**Figure 5 Lower Panel**). Although carbon starvation was unable to rescue *ERG110* expression levels in the absence of elF2α (**Supplemental Figure 6**), it completely restored the expression of *TRR1* (**Figure 6C**). These, along with the complementary experiments performed in Figure 4 and Figure 5, strongly suggest that rapid but transcript specific translational inhibition in response to H_2_O_2_ is necessary for the expression of critical oxidative stress response transcripts. To determine if the restored transcript kinetics translated to increased oxidative stress resistance in the *gcn2*Δ strain, cultures were treated with a Live-Dead stain following glucose starvation and exposure to H_2_O_2_ (**Figure 6D Schematic**). The percentage of dead yeast was determined by flow cytometry and quantified as a percentage of the total. Removing glucose just one hour prior to the addition of 2mM H_2_O_2_ returned the oxidative stress resistance of the *gcn2*Δ strain to Wild Type levels (**Figure 6D**).

To see if the failure of *TRR1* transcript expression in the *gcn2*Δ strain was due to a defect in the translational expression of the respective transcription factor, *ATF1*, we performed northern blot analysis against sucrose gradient fractionated RNA. There were no observed differences in *ATF1* distribution in the polysomes between the two strains (**Figure 5A Middle Panel**). Furthermore, the distribution of *ATF1* upon carbon starvation and hydrogen peroxide exposure was not different between the strains (**Figure 5A Bottom Panel**).

## Discussion

The extent and severity of translational repression in response to H_2_O_2_ derived oxidative stress in *C. neoformans* is driven largely by the phosphorylation of eIF2α. This repression is not absolute, however, as puromycin incorporation is still detected even after being exposed to high levels of H_2_O_2_ (**Figure 1C**). Therefore, it seems that a subset of transcripts may possess elements that allow them to be translated under conditions that limit the active ternary complex. Previous examples of mRNAs possessing uORF being translationally favored under conditions of elF2α phosphorylation are known in other systems (21, 23). 122 predicted uORFs are found to be conserved across four sequenced *Cryptococcus* strains and may represent a major post-transcriptional regulatory strategy for the expression of these transcripts (25). Furthermore, recent (pre-print) ribosome profiling results indicate that over a third of *C. neoformans* transcripts possess uORF that affect translation (33). Our results suggest that certain stressors that activate Gcn2 may translationally favor the expression of the annotated ORF of these transcripts in *C. neoformans*.

Failure to phosphorylate eIF2α resulted in the absence of translational inhibition and the increased stability of *RPL2*. This stability is recapitulated in the wild type strain upon treatment with cycloheximide, suggesting that ribosome association protects the mRNA from decay factors (**Figure 4B**). Promoting translational suppression through glucose starvation in *gcn2*Δ was able to rescue *TRR1* expression and oxidative stress resistance (**Figure 6C-D**). How translational suppression restored expression of *TRR1* still remains to be determined. Based on the abundance of the transcription factor *ATF1* and its high polysome association, one would assume that the expression of *TRR1* would be higher than expected, yet the opposite exists. Interestingly, hydrogen peroxide stress has been found to cause a buildup in protein aggregates believed to be caused by protein misfolding (34). Loss of mRNA surveillance pathways that accelerate the decay of certain transcripts further exacerbates protein aggregate formation (35). Could translational suppression favor the proper folding of the transcription factor and is protein misfolding the true cause of death in *C. neoformans* exposed to H_2_O_2_? It is an interesting hypothesis given the recent appreciation of co-translational protein folding (36).

Altogether, our results suggest a tight interconnectedness between translation and mRNA decay that drastically affects the ability of the fungal pathogen to adapt to oxidative stress. Disrupting the translational response to H_2_O_2_ resulted in changes in the stress response at the transcript level indicating the yet unappreciated role ribosome availability plays in regulating mRNA levels and ultimately their expression during stress. The scientific community’s understanding of Eukaryotic gene regulation has largely been in the context of steady state exponential phase growth conditions. Our results suggest that when this context changes, so do the dynamics of gene regulation.

## Acknowledgments

We would like to thank all members of the Panepinto Lab for discussion relating to the project’s design and implementation.

---

**Supplemental Figure 1.**
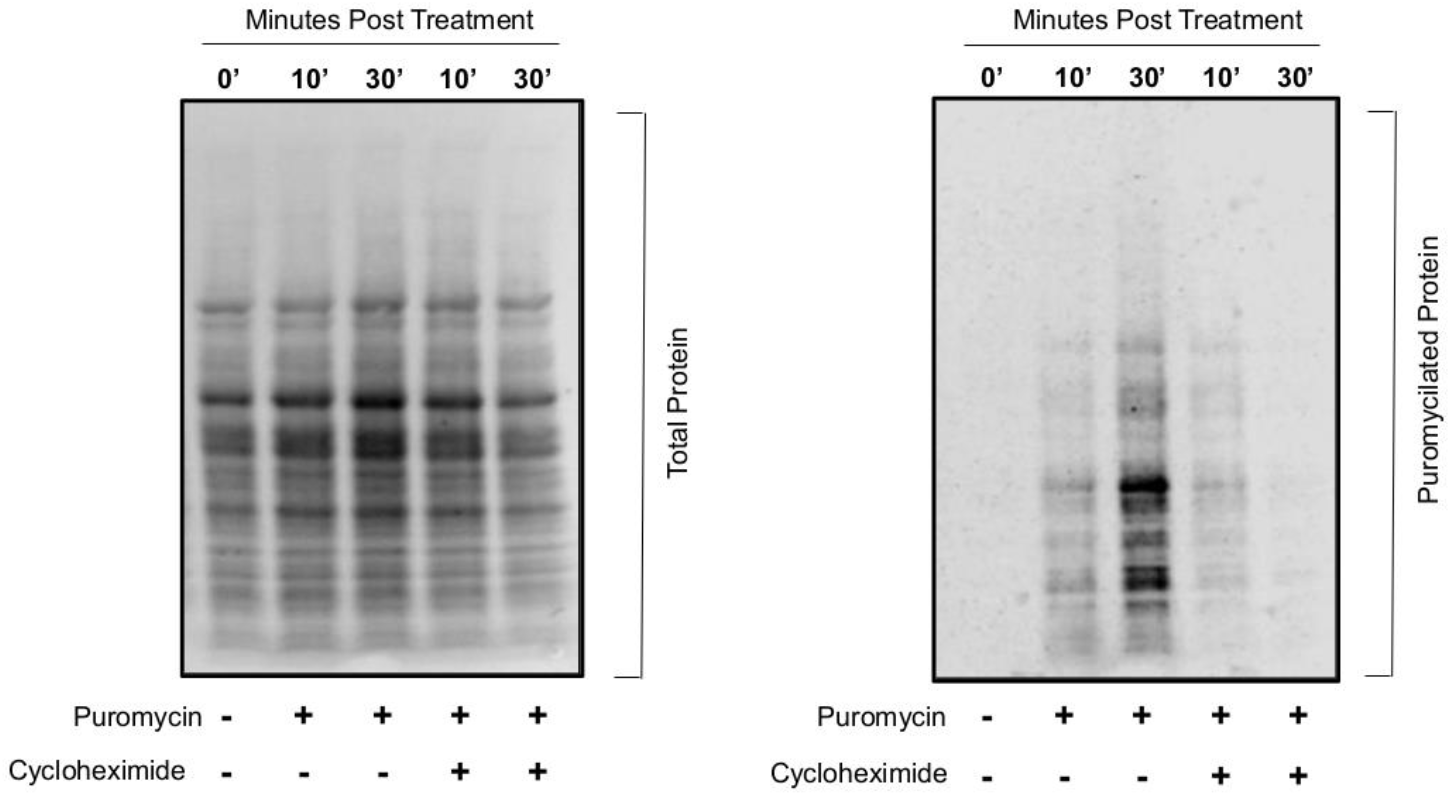
Experimental validation of the use of puromycin in C. neoformans to assess translational output. Yeast were grown to midlog in minimal media supplemented with 2% dextrose. At that time cultures were harvested in the presence of either puromycin, cycloheximide or both for the indicated amount of time. The image on the left represents total protein loaded into each lane following membrane transfer. Signal intensity on the right is derived from immunoblotting against nascent proteins covalently bound with puromycin during translation. Detection of non-specific binding of the α-puromycin antibody is low (Lane 1). Cyclohexime was used as a negative control to assess the need for translational elongation for puromycin incorporation.

**Supplemental Figure 2.**
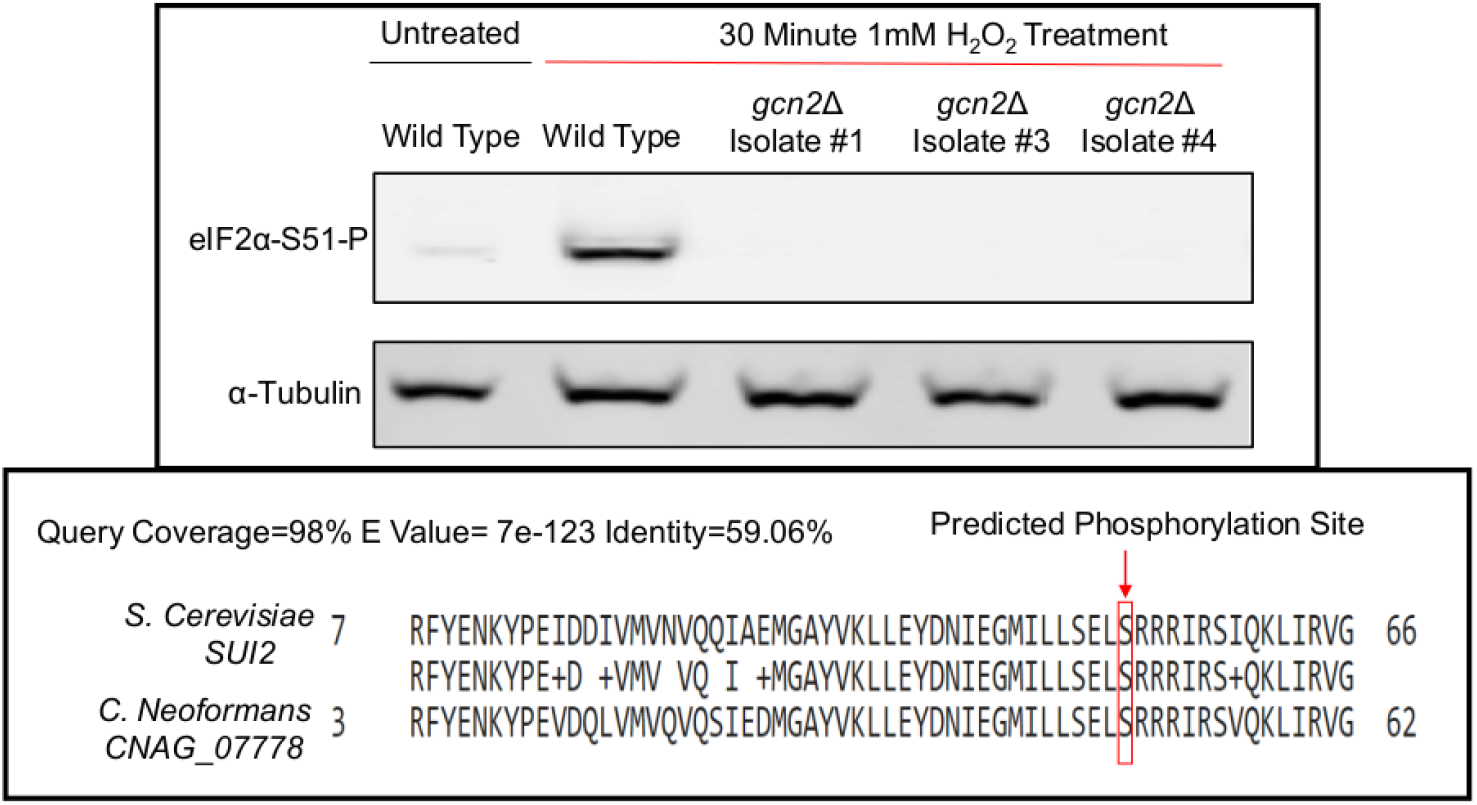
Gcn2 is the sole kinase of eIF2α in C. neoformans. Yeast were grown to midlog in YPD. Cultures were then exposed to 1mM H_2_O_2_ for 30 minutes to induce eIF2α phosphorylation. Western blot analysis was performed using an antibody that recognizes the phosphorylated form of eIF2α. Bottom panel represents the homology between the N-terminus of eIF2α in *C. neoformans* and *S. cerevisiae*, with the targeted serine outlined in red. 92% identify exists between the first 100 amino acids.

**Supplemental Figure 3.**
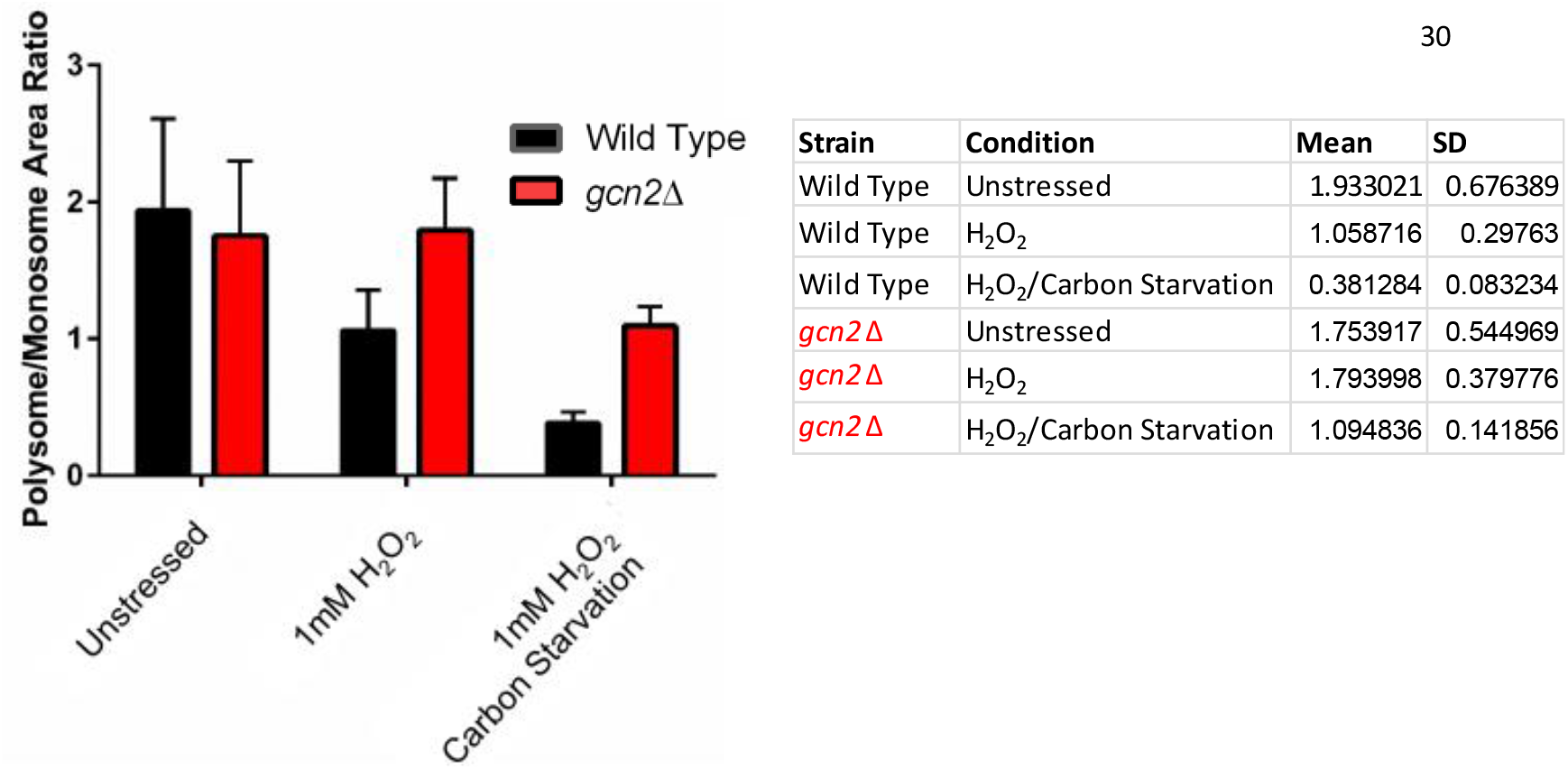
Hydrogen peroxide and carbon starvation results in translational suppression. Yeast were grown to midlog in minimal media supplemented with 2% dextrose. Cultures were then pelleted by centrifugation and were either resuspended in minimal media with or without 2% dextrose in the presence of 1mM H_2_O_2_. Cultures were harvested after 30 minutes and polysome profiles were obtained. Following trace acquisition, the area under the curves corresponding to the polysome and monosome peaks were quantified. Extent of translational suppression was determined by comparing the total polysome peak area to that of the monosome peak area. Error bars represent SD of three biological replicates, with exact numercal values depicted on the right side panel. One-Way nonparametric analysis (Kruskal-Wallis test) determined the results significant with Wild Type p= 0.0036 *gcn2*Δ = 0.05.

**Supplemental Figure 4.**
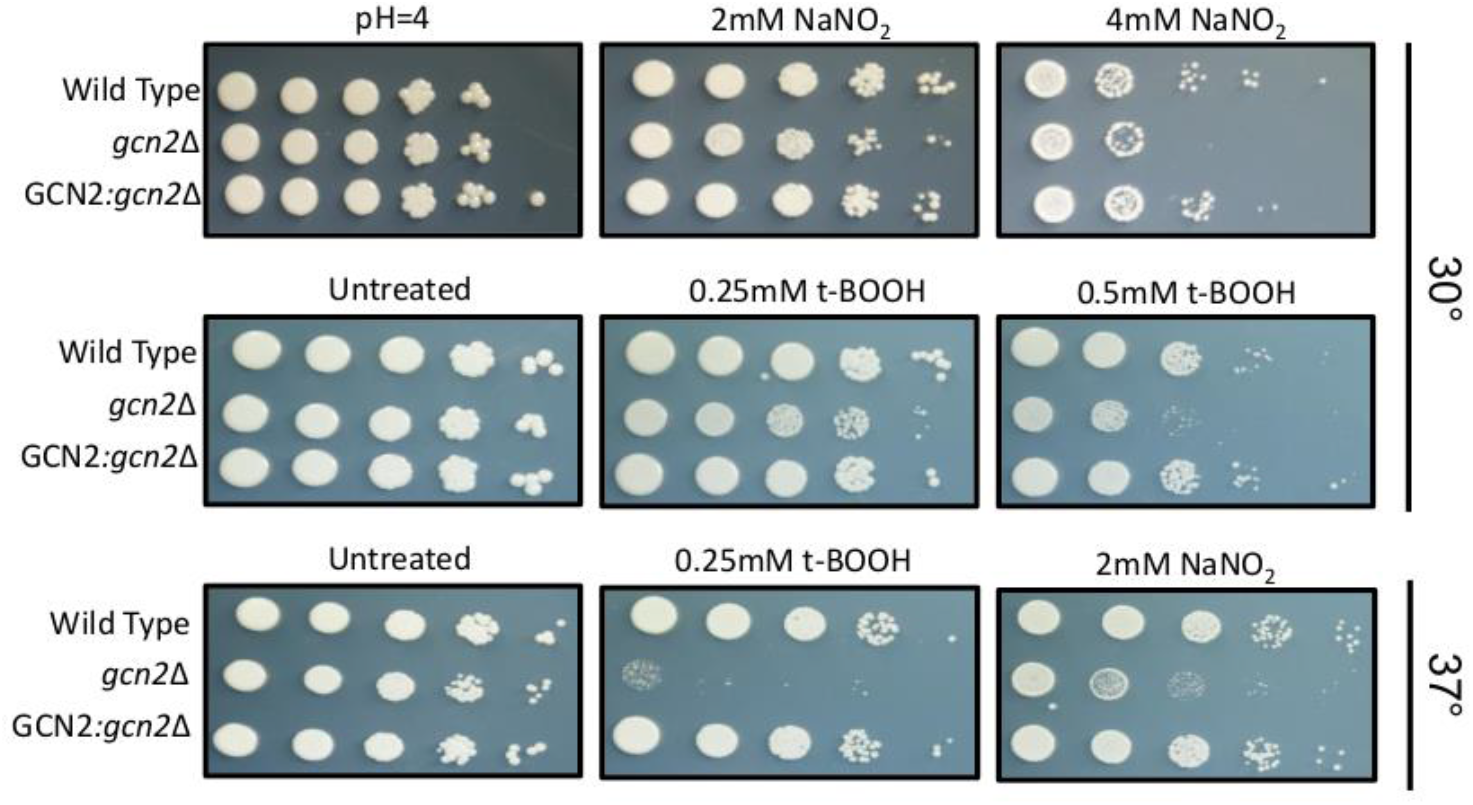
Gcn2 is required for growth in the presence of both RNS and ROS. Strains were grown in YPD for 16 hours prior to suspension in water to an OD_600_ of 1.0. Cultures were serial diluted 10 fold onto agar plates containing minimal media and 2% dextrose imbued with the indicated stressor. Tert-butyl hydroperoxide (t-BOOH) generates ROS where else sodium nitrite (NaNO_2_) when incubated at a low pH dissociates to NO^−^, which generates RNS. Plates where incubated at either 30° or 37° for three days.

**Supplemental Figure 5.**
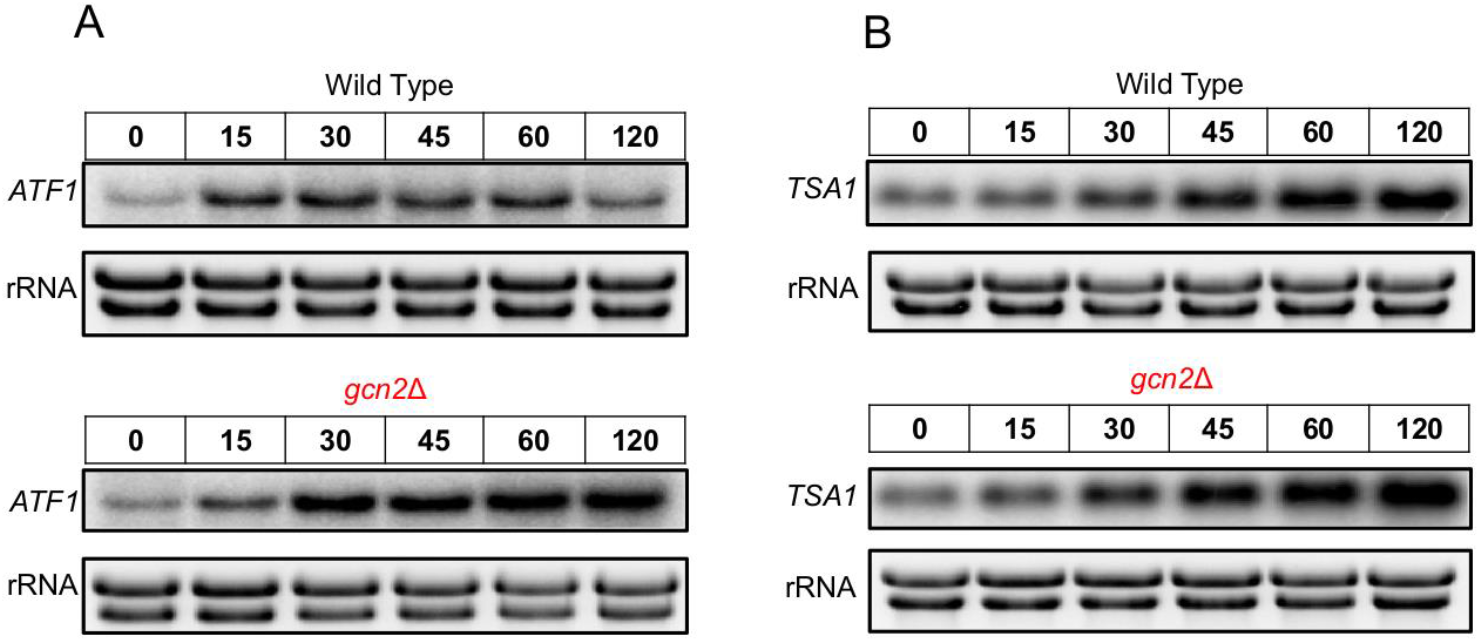
Expression of transcripts associated with TRR1 transcript induction is still intact in the absence of Gcn2. Cultures were grown to exponential phase in YPD and where subjected to 1mM H_2_O_2_. Aliquots where harvested at indicated time points during incubation, whole RNA was extracted, and northern blot assays were performed probing for **(A)** *ATF1* and **(B)** *TSA1*.

**Supplemental Figure 6.**
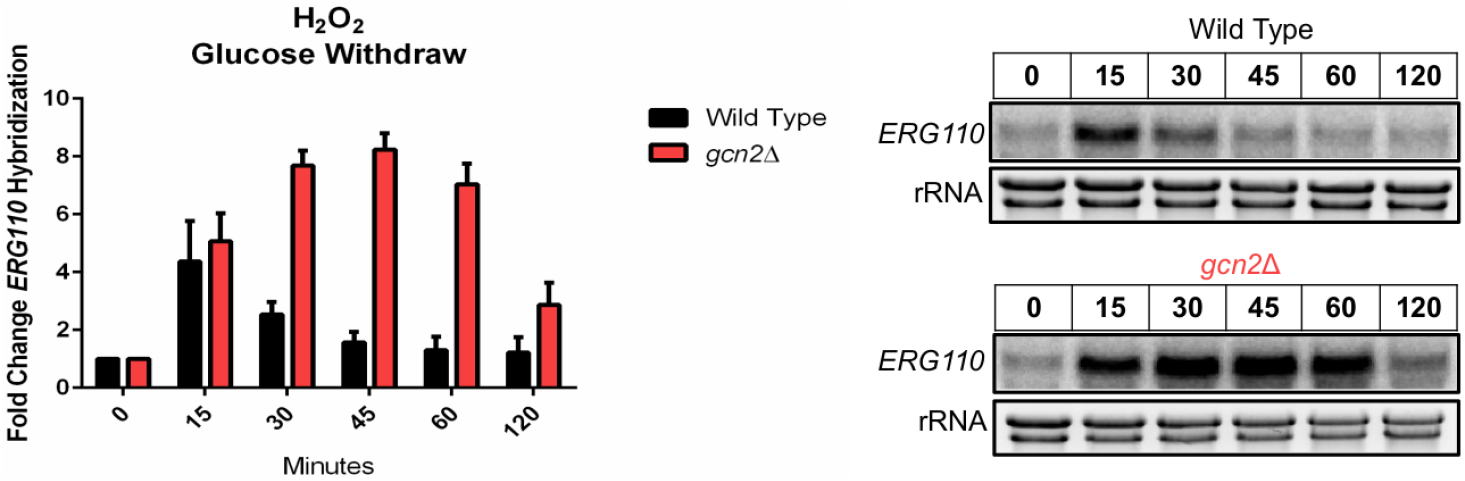
Glucose starvation fails to suppress H_2_O_2_ induced ERG110 levels in the absence of Gcn2. Cultures were grown to midlog in minimal media supplemented with 2% dextrose. Cultures were then pelleted, washed with water, and the re-suspended in carbonless minimal media with 1mM H_2_O_2_. Aliquotes were harvested and total RNA was isolated at indicated time points. *ERG110* levels probed for by northern blot analysis. n=2

